# Intracranial functional haemodynamic relationships in patients with cerebral small vessel disease

**DOI:** 10.1101/572818

**Authors:** Gordon W Blair, Michael J Thrippleton, Yulu Shi, Iona Hamilton, Michael Stringer, Francesca Chappell, David Alexander Dickie, Peter Andrews, Ian Marshall, Fergus N Doubal, Joanna M Wardlaw

## Abstract

**Background:** Cerebral small vessel disease (SVD) is a major cause of stroke and dementia. The underlying cerebrovascular dysfunction is poorly understood. We investigated cerebrovascular reactivity, blood flow, vascular and cerebrospinal fluid (CSF) pulsatility, and their independent relationship to SVD features, in patients with minor ischaemic stroke and MRI evidence of SVD.

**Methods:** We recruited patients with minor ischaemic stroke and assessed CVR using Blood Oxygen Level Dependent (BOLD) MRI during a hypercapnic challenge, cerebral blood flow, vascular and CSF pulsatility using phase contrast MRI, and structural MR brain imaging to quantify white matter hyperintensities (WMH) and perivascular spaces (PVS). We quantified CVR in seven white matter and six subcortical grey matter regions, measured blood flow in carotid and vertebral arteries, intracranial venous sinuses, internal jugular veins and CSF flow at the aqueduct and foramen magnum. We used multiple regression to identify SVD features, blood flow and pulsatility parameters associated with CVR, controlling for patient characteristics.

**Results:** In 53 of 60 patients with complete data (age 68.0±8.8, 74% male, 75% hypertensive), CVR in grey and white matter decreased with increasing blood pressure (BP, respectively −0.001%/mmHg, p=0.01 and −0.006%/mmHg, p=0.01, per mmHg increase in BP). After controlling for age, gender and systolic BP, white matter CVR decreased with increasing WMF volume (−0.01%/mmHg per log10 increase in WMH volume, p=0.02) and basal ganglia PVS (−0.01%/mmHg per point increase in PVS score, p=0.02). White matter CVR decreased with increasing venous pulsatility (superior sagittal sinus −0.03%/mmHg, p=0.02, per unit increase in pulsatility index) but not with cerebral blood flow (p=0.58). Lower foramen magnum CSF stroke volume was associated with worse white matter CVR (0.04%/mmHg per ml increase in stroke volume, p=0.04) and increased basal ganglia PVS.

**Conclusions:** Contemporaneous assessment of CVR, intracranial vascular and CSF pulsatility demonstrates important interrelationships of these vascular functions in humans. Decreased CVR, increased venous pulsatility and reduced foramen magnum CSF stroke volume suggests that dynamic vascular dysfunctions underpin PVS dysfunction and WMH development. Improved understanding of microvascular dysfunction and CSF dynamics offers new intervention targets to reduce SVD lesion development and their impact on cognitive dysfunction and stroke.

## Introduction

Cerebral small vessel disease (SVD) causes lacunar ischaemic stroke, vascular cognitive impairment and dementia, intracerebral haemorrhage and gait and bladder dysfunction with prevalence increasing with age.(1) Diagnosis is via acute presentation with a lacunar stroke syndrome, cognitive dysfunction or mobility problems, and with neuroimaging findings such as a recent small subcortical infarct, white matter hyperintensities (WMH), lacunes or microbleeds.(2)

There is no specific treatment for SVD as yet.(3) An improved understanding of disease mechanisms is required to identify drug targets. The role of cerebral blood flow (CBF) in SVD is complex: whilst lower CBF is associated with more severe SVD in cross-sectional studies, most longitudinal studies have not found lower CBF to precede development or progression of SVD,(4) and dynamic measures of CBF may be more sensitive to small vessel dysfunction. Reliable measures of vascular function would help screen potential treatments in early phase trials(5) prior to definitive but costly trials with clinical endpoints that require large sample sizes and long follow up periods.(6) Sensitive measures of early cerebrovascular dysfunction might also help personalise therapies.

Cerebrovascular reactivity (CVR) measures the ability of cerebral blood vessels to increase CBF in response to metabolic demands.(7) CVR can be assessed by measuring blood flow changes in response to a vasoactive stimulus such as increased carbon dioxide in inspired air, breath holding, or acetazolamide injection.(8) Transcranial Doppler (TCD) ultrasound is commonly used but only measures large basal cerebral arteries and lacks spatial resolution.(9,10) Magnetic resonance imaging (MRI) techniques measure CVR in all brain tissues, including the subcortical areas commonly affected by SVD.(11) So far, MRI-based CVR has been assessed in relatively few small studies in SVD, but used different CVR challenges,(12-16) and did not account for clinically-relevant variables (e.g. age, blood pressure [BP], gender) that could confound CVR associations with SVD features.(12)

Vascular pulsatility measured outside the brain is associated with increasing WMH(17,18) consistent with the hypothesis that SVD is associated with increased arterial stiffening. Also, middle cerebral artery (MCA) pulsatility increases with increasing WMH on TCD,(9,10) but TCD cannot measure intracranial veins, venous sinuses, or cerebrospinal fluid (CSF) pulsatility, precluding a complete assessment of intracranial pulsatility across key fluid compartments. MRI with phase contrast angiography can assess pulsatility and flow in the intracranial arteries, veins, venous sinuses and CSF spaces (e.g. aqueduct, foramen magnum) within one examination, concurrently with CVR, making it possible to assess relationships between CVR, CBF and pulsatility directly. However, there are very few studies of intracranial pulsatility assessed with MRI in ageing or SVD(19) and no studies in humans of CVR assessed concurrently with intracranial vascular pulsatility, CSF flow dynamics, or resting CBF, to provide a complete assessment of intracranial haemodynamics.

We aimed to assess CVR and patient demographic variables, vascular risk factors and imaging features of SVD, concurrently with total CBF, arterial, venous and CSF pulsatility. We studied independent patients with a past history of minor ischaemic stroke stratified by SVD burden as representative of a high risk group for the clinical effects of SVD. We and hypothesised that more severe SVD features on neuroimaging would be associated with reduced CVR, that reduced CVR would associate with increased vascular pulsatility, and that these relationships would persist after controlling for patient demographic and vascular risk factors.

## Methods

### Patients

We recruited patients presenting to our regional stroke service between October 2014 and April 2016 with symptomatic non-disabling ischaemic stroke (Modified Rankin score ≤3). We also invited such patients who had participated in recent previous prospective studies.(20) All patients had a clinical stroke diagnosis confirmed by a specialist stroke physician and brain imaging that either confirmed a relevant recent infarct or, if no recent infarct was visible, excluded any other cause of the presenting symptoms.(20)

We excluded patients who were pregnant, unable to lie flat, had contraindications to MRI (including claustrophobia), moderate to severe chronic respiratory disease or symptomatic cardiac failure, personal history or first degree relative with subarachnoid haemorrhage or intracranial aneurysm, uncontrolled hypertension, or atrial fibrillation with fast ventricular response.

All participants provided written informed consent prior to enrolment. The study was approved by the UK Health Research Authority National Research Ethics Service Committee East Midlands, Nottingham 1 (ref. 14/EM/1126).

We performed the study assessments at least one month after the patient’s stroke to prevent haemodynamic changes in the acute phase of stroke (both stroke related and due to commencing vasoactive secondary prevention medications) from interfering with the interpretation of vascular function. We requested participants not to consume caffeinated drinks on the day of assessment.

We recorded detailed medical histories including clinical characteristics of presenting stroke, vascular risk factors, concurrent medications, and investigations performed for the presenting stroke (diagnostic brain MRI, carotid ultrasound imaging, haematology, biochemistry, electrocardiograms).

We classified the clinical stroke syndrome according to the Oxford Community Stroke Project classification(21) (independently assessed by two stroke physicians, FDand GB) and the lesion type seen on imaging (independently assessed by JMW and YS). Where no recent ischaemic lesion was evident, the final stroke type was assigned using the clinical classification. Where the imaging lesion was discordant with the clinical stroke type, the imaging type was used as the final classification, as previously.(22)

We recorded blood pressure (BP) seven times at standard time points before, during and after MRI.

### MRI

We performed brain MRI using a 1.5 Tesla GE research MRI scanner (Signa H Dxt, General Electric, Milwaukee, WI) operating in research mode and an 8-channel phased array head coil. Total imaging time was circa 75 minutes for each participant, but patients were allowed to move and have a natural break between each section of scans to ensure they remained comfortable. We acquired 3D T1-weighted, axial T2-weighted, fluid attenuated inversion recovery (FLAIR) and gradient echo sequences. We performed BOLD MRI scanning with CO_2_ challenge at 4mm isotropic resolution acquiring a whole brain volume every three seconds.(11) We performed phase contrast MRI to measure pulsatility in the internal carotid arteries, intracranial venous sinuses, CSF flow (aqueduct, foramen magnum) and total CBF, as described in detail.(23) Full parameters of all MR sequences are in Supplementary Table 1.

**Table 1:**
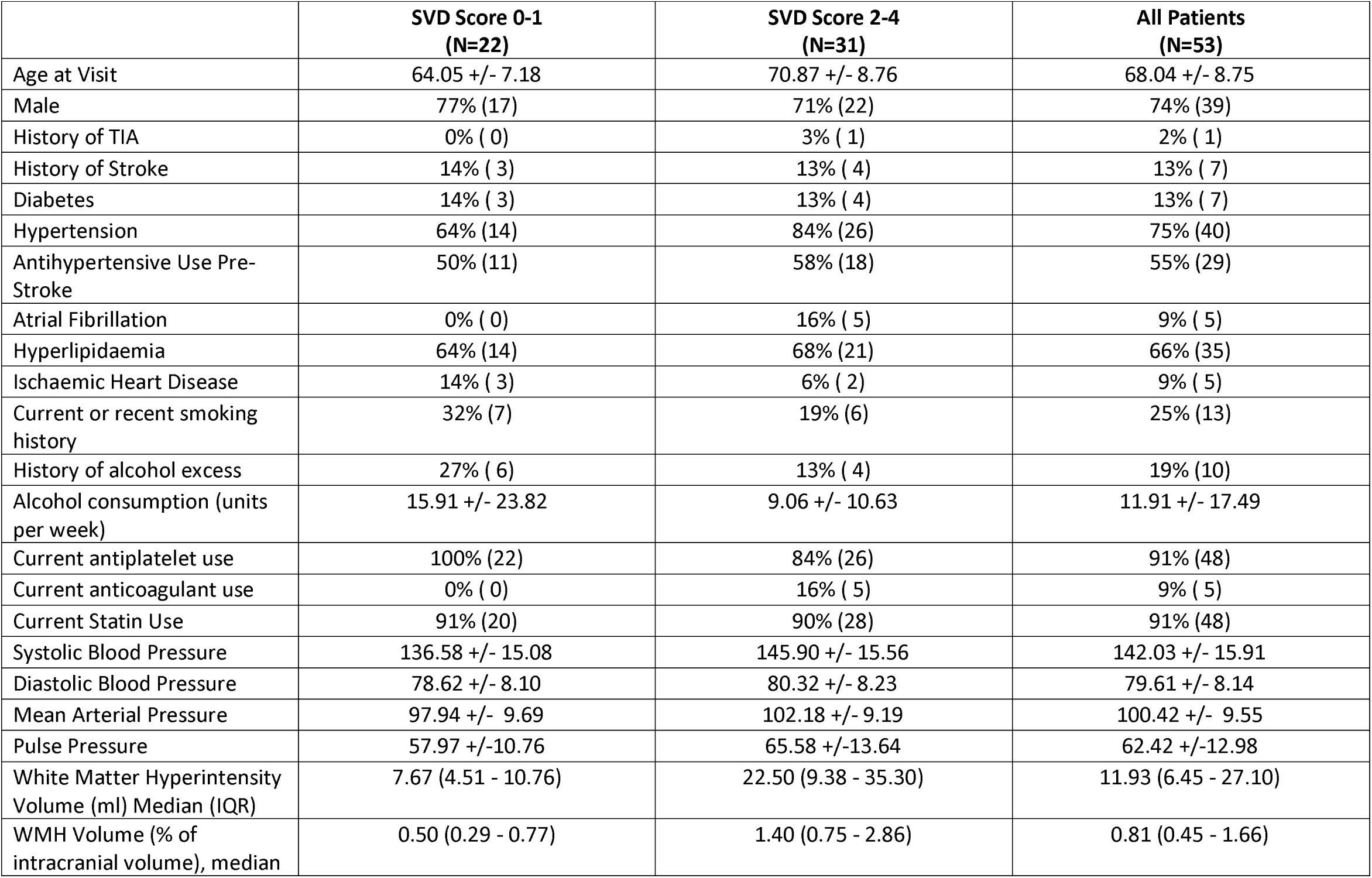
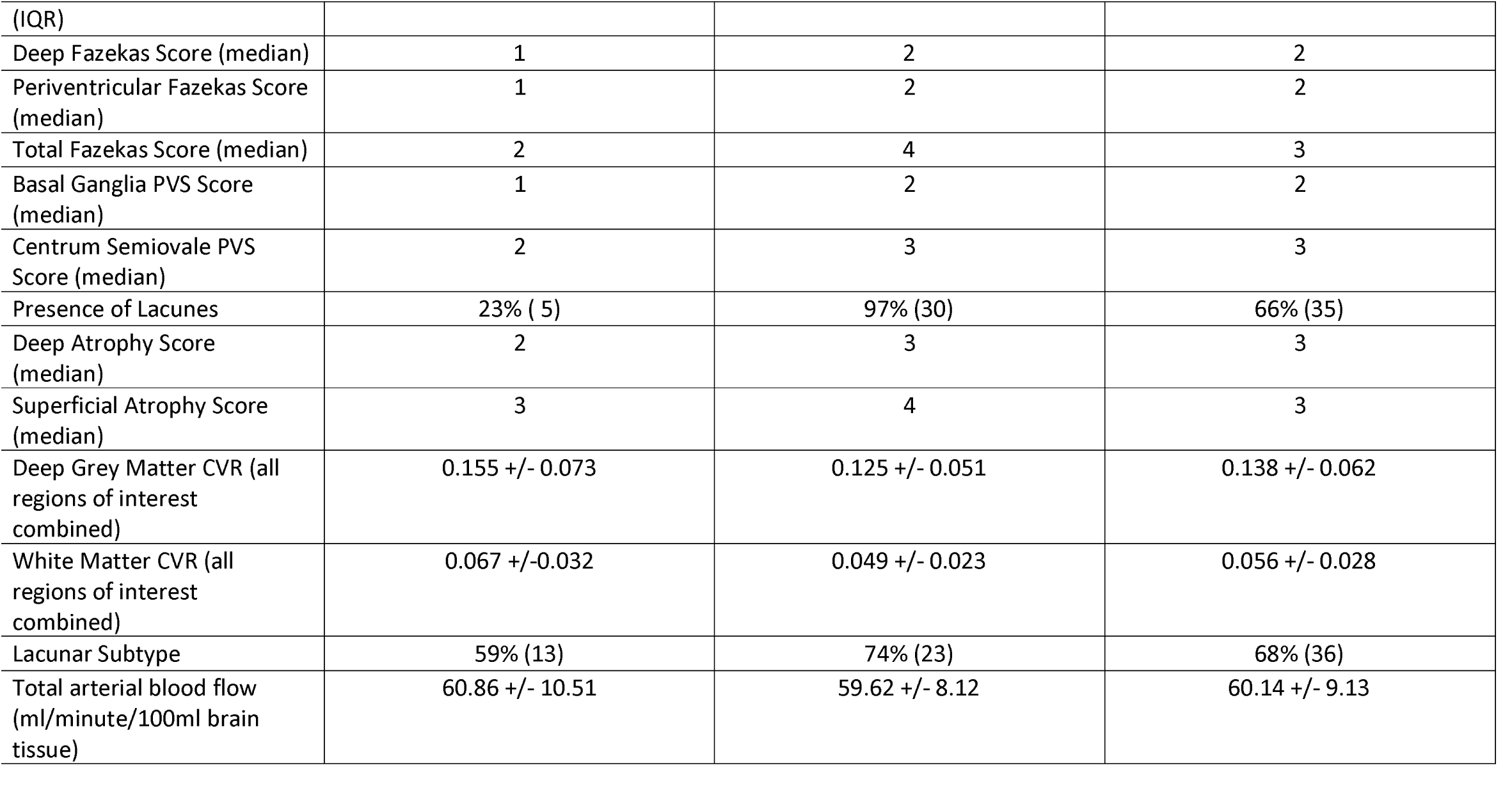
Demographic data of participants by SVD score. Continuous variables expressed as mean +/-SD. Categorical variables expressed as n (%).

The CVR method used BOLD 2-dimensional echo-planar imaging with a CO_2_ challenge (detailed in(11)). Briefly, during MRI, patients wore an anaesthetic face mask, carefully fitted to avoid gas leak between mask and face, attached to a bespoke unidirectional breathing circuit (Intersurgical, Wokingham, UK). Monitoring equipment recorded pulse rate, oxygen saturation, BP (Millennia 3155A and Magnitude 3150 MRI; Invivo, Best, The Netherlands) and end-tidal CO_2_ (EtCO2; CD3-A AEI Technologies, Pittsburgh, USA) throughout the examination. During a 12-minute BOLD MRI scan, patients breathed medical air and 6% CO_2_ in air (BOC Special Products, UK) alternately, delivered as two minutes air, three minutes CO_2_, two minutes air, three minutes CO_2_, finishing with two minutes air. We instructed patients to expect a change in smell and breathing pattern (deeper, faster or more forceful breathing), then each patient tried the facemask and gasses before entering the scanner room. Patients were instructed to breathe normally and to press a buzzer to stop the scan if required. Scanning commenced when the patent was fully positioned in the scanner bore, indicated they felt comfortable, and blood pressure and heart rate readings indicated no sign of anxiety or distress. A physician monitored the patient during CO_2_ inhalation.

Arterial and venous flow waveforms were measured as described previously. (23-25) Briefly, we used a 2D cine phase-contrast sequence with retrospective peripheral pulse gating to acquire 32 velocity images per cardiac cycle. We used the following slice locations to measure flow in the different structures: a slice superior to the carotid bifurcation and perpendicular to the internal carotid artery (ICA) walls to measure flow in the ICA, vertebral arteries (VA) and internal jugular veins (IJV); a coronal-oblique slice intersecting the superior sagittal sinus approximately 2cm above the torcular and through the midpoint of the straight sinus to measure superior sagittal sinus (SSS), straight sinus (StS) and transverse sinus (TS) flow; a slice perpendicular to the aqueduct for aqueduct CSF flow and an axial slice at the cranio-cervical junction for foramen magnum CSF flow.

### Image Processing and Analysis

Investigators were blinded to patient clinical data, CVR, pulsatility and CBF results for all analysis.

Structural image analysis of SVD features was performed according to the STRIVE criteria(2) under the supervision of an expert neuroradiologist (JMW).

We (YS, JMW) scored the following features: WMH using the Fazekas scale, summing periventricular and deep WMH scores to give a score from 0-6; perivascular spaces (PVS), scored separately in the basal ganglia and centrum semiovale, using a validated, semi-quantitative ordinal scale (range 0-4); lacunes (location, number); and microbleeds (BOMBS scale) presence/absence and total number(2); and total SVD score (0-4) by combining WMH, lacunes, microbleeds and PVS scores as described previously.(26)

We co-registered each subject’s structural images. We calculated WMH volumes using a validated semi-quantitative technique described previously.(23) Briefly we generated WMH probability maps for each subject using FLAIR and T1W image data. Hyperintense outliers within the white matter surface were defined as voxels with a z score ≥ 1.5 on FLAIR to create an initial estimate of WMH volume. We produced final estimates using 3D smoothing to account for partial volume effects and reduce noise, before manually removing the index and any prior stroke lesions. We segmented normal appearing tissues (cortical grey matter, subcortical grey matter, white matter, cerebellum) and whole brain volume from each subject’s T1W data and local population specific probability maps.(27) We calculated intracranial volumes using a semi-automatic method based on T2*W images. All tissue masks were visually inspected and manually corrected as necessary.

We processed CVR images as described previously.(11) We generated voxel-wise CVR maps by regressing the BOLD signal against EtCO2 and the BOLD scan number (to account for signal drift). CVR is expressed as % BOLD signal change/mmHg change in EtCO2, based on the EtCO2 regressor in the model. We made additional adjustment for the delay time between BOLD signal change and EtCO2 change to minimise the residual sum of squares individually for each voxel, further adjusted by 4 seconds to account for the delay between exhalation and detection on the EtCO2 monitor caused by the eight metre sample line.

We realigned BOLD images (using SPM 8, https://www.fil.ion.ucl.ac.uk/spm/software/spm8/) prior to determining the transformation between BOLD and T2W image spaces (using FSL FLIRT(28)). We then manually drew three subcortical grey matter (thalamus, putamen, caudate head) and four subcortical white matter (frontal, posterior, periventricular and centrum semiovale) regions of interest (ROI) on T1W images before transfer to the BOLD images. Voxels that were part of large vessels or the patient’s stroke lesion were manually excluded. We extracted the mean signal across the ROI and fitted the CVR model to that data. We also averaged all grey and white matter regions to give a combined deep grey matter and white matter CVR value.

To process the phase contrast data we drew manual ROI’s around right and left ICAs and VAs, the sagittal, straight, right and left transverse sinuses and internal jugular veins, the aqueduct and the foramen magnum subarachnoid space. Background ROIs were placed close to the ROI’s to correct background phase error by subtracting the background velocity (’noise’) from the ROI velocity. We calculated sum flow and mean velocity for bilateral structures, then total CBF as the sum of ICA and VA flow, normalised to total brain volume and expressed as ml/min/100ml brain tissue. Pulsatility index (PI) in each structure was calculated as (Flow_maximum_ – Flow_minimum_)/Flow_mean_; Resistivity index (RI) was calculated as (Flow_maximum_ – Flow_minimum_)/Flow_maximum_. The reproducibility of this approach is published.(23)

### Statistical Analysis

We performed statistical analyses using R version 3.3.0 (https://cran.r-proiect.org/) and the additional packages Hmisc, texreg, data.table, htmlTable, car and psych. We assessed distribution of all variables prior to analysis and log transformed the WMH volumes and atrophy scores due to a skewed distribution. The BP values presented are means of the seven readings taken across the visit. We assessed the relation of demographic factors, SVD features, pulsatility and CBF parameters to CVR values using univariate and then multiple linear regression, adjusting for age, systolic BP and gender, based on those factors identified in our systematic review as potentially influencing CVR,(12) (univariate associations are presented only for completeness). Additionally we adjusted for WMH volume in some models to assess if relationships were independent of a co-association, since we found previously that failure to control for WMH was a potential confound in assessments of CVR in patients with SVD.(6) We examined the normality of residuals (QQ plots and histograms) and heteroscedascity (residual versus fitted values) to assess modelling assumptions. We checked variable inflation factors to assess for collinearity between variables, with a limit of two applied.(29)

## Results

### Study Population

We recruited 60 patients; 53 subjects completed the full CVR scanning protocol with complete and fully analysable data. Three withdrew due to claustrophobia when wearing the facemask in the MRI scanner, in one movement artefact precluded analysis, two failed to show any EtCO_2_ or BOLD signal change with CO_2_ (likely due to a poor fitting mask), and in one patient the acquisition was stopped after initial structural scanning showed an incidental asymptomatic subdural haematoma.

The cohort had a mean age of 68.0±8.8 years (range 52-88 years), 39 (73%) were male. Median NIHSS at presentation with stroke was 2 (range 0-5). At the time of CVR scanning, 75% had hypertension, 66% had hyperlipidaemia, 13% had diabetes, 30% were current smokers or had stopped within the past 12 months and 19% consumed alcohol regularly in excess of recommended limits (table 1). Subjects were scanned at a median of 92 days post stroke (range 32-1768 days).

Imaging features of SVD were common: 27 patients (51%) had deep Fazekas score ≥2 and 29 (55%) had periventricular Fazekas ≥2 indicating moderate to severe WMH; 17 (32%) had basal ganglia PVS score ≥3 and 28 (53%) had centrum semiovale PVS score ≥3 indicating high PVS visibility; 35 (66%) had lacunes; and four patients (8%) had microbleeds.

### Univariate Analysis

#### CVR and Patient characteristics

Higher systolic BP and pulse pressures were associated with reduced CVR in white matter (all p=0.01) and grey matter (all p=0.01; Supplementary Table 2). Reduction of white matter CVR with increasing age did not reach statistical significance (p=0.09). We found no other statistically significant associations with reduced CVR.

**Table 2:** CVR association with stroke subtype and SVD imaging markers. All multivariate regression models adjusted for age, sex and systolic blood pressure. Standardised ß co-efficient 95% confidence interval and p value.

|  | **Deep Grey Matter CVR** | **White Matter CVR** | **VIF Range** |
| --- | --- | --- | --- |
| Stroke Type (lacunar vs Cortical) | $\beta = 0.006$<br>(-0.031, 0.043)<br>$p = 0.73$ | $\beta = 0.001$<br>(-0.015, 0.264)<br>$p = 0.89$ | 1.07-1.16 |
| WMH Volume (normalised to ICV) | $\beta = -0.021$<br>(-0.043, 0.000)<br>$p = 0.05$ | $\beta = -0.012$<br>(-0.021, -0.002)<br>$p = 0.02$ | 1.10-1.58 |
| Periventricular Fazekas Score | $\beta = -0.023$<br>(-0.047, 0.002)<br>$p = 0.07$ | $\beta = -0.013$<br>(-0.023, -0.002)<br>$p = 0.02$ | 1.12-1.48 |
| Deep Fazekas Score | $\beta = -0.011$<br>(-0.033, 0.011)<br>$p = 0.30$ | $\beta = -0.005$<br>(-0.015, 0.005)<br>$p = 0.30$ | 1.10-1.35 |
| Total Fazekas | $\beta = -0.009$<br>(-0.021, 0.003)<br>$p = 0.14$ | $\beta = -0.005$<br>(-0.010, 0.001)<br>$p = 0.08$ | 1.11-1.41 |
| PVS Score – Basal Ganglia | $\beta = -0.019$<br>(-0.039, 0.001)<br>$p = 0.07$ | $\beta = -0.011$<br>(-0.019, -0.002)<br>$p = 0.02$ | 1.10-1.41 |
| PVS Score – Centrum semiovale | $\beta = -0.017$<br>(-0.034, 0.001)<br>$p = 0.06$ | $\beta = -0.007$<br>(-0.014, 0.001)<br>$p = 0.08$ | 1.11-1.21 |
| Lacunes | $\beta = -0.017$<br>(-0.056, 0.022)<br>$p = 0.38$ | $\beta = -0.010$<br>(-0.028, 0.007)<br>$p = 0.23$ | 1.11-1.29 |
| Deep Atrophy | $\beta = -0.011$<br>(-0.046, 0.024)<br>$p = 0.53$ | $\beta = -0.010$<br>(-0.025, 0.005)<br>$p = 0.20$ | 1.10-1.33 |
| Superficial Atrophy | $\beta = -0.011$<br>(-0.054, 0.032)<br>$p = 0.62$ | $\beta = -0.006$<br>(-0.024, 0.013)<br>$p = 0.56$ | 1.12-1.36 |
| Combined Atrophy | $\beta = -0.012$<br>(-0.055, 0.030)<br>$p = 0.56$ | $\beta = -0.009$<br>(-0.028, 0.009)<br>$p = 0.32$ | 1.10-1.40 |
| Brain Parenchymal<br>Volume (normalised to<br>ICV) | $\beta = 0.38$<br>(-0.175, 0.936)<br>$p = 0.17$ | $\beta = 0.010$<br>(-0.240, 0.259)<br>$p = 0.94$ | 1.12-1.32 |
| SVD Score (High vs Low) | $\beta = -0.026$<br>(-0.063, 0.011)<br>$p = 0.17$ | $\beta = -0.011$<br>(-0.028, 0.005)<br>$p = 0.17$ | 1.14-1.57 |

WMH = white matter hyperintensity, PVS = perivascular space, ICV = intracranial volume, VIF = variable inflation factor

#### CVR and SVD Features

Numerous SVD features were associated with lower white matter CVR (Figure 1, Supplementary Table 2): increased WMH volume, worse WMH Fazekas scores, more PVS, presence of lacunes, deep atrophy and total SVD score (all p<0.05). Reduced grey matter CVR was only associated with periventricular WMH Fazekas score and basal ganglia PVS (both p<0.05).

**Figure 1:**
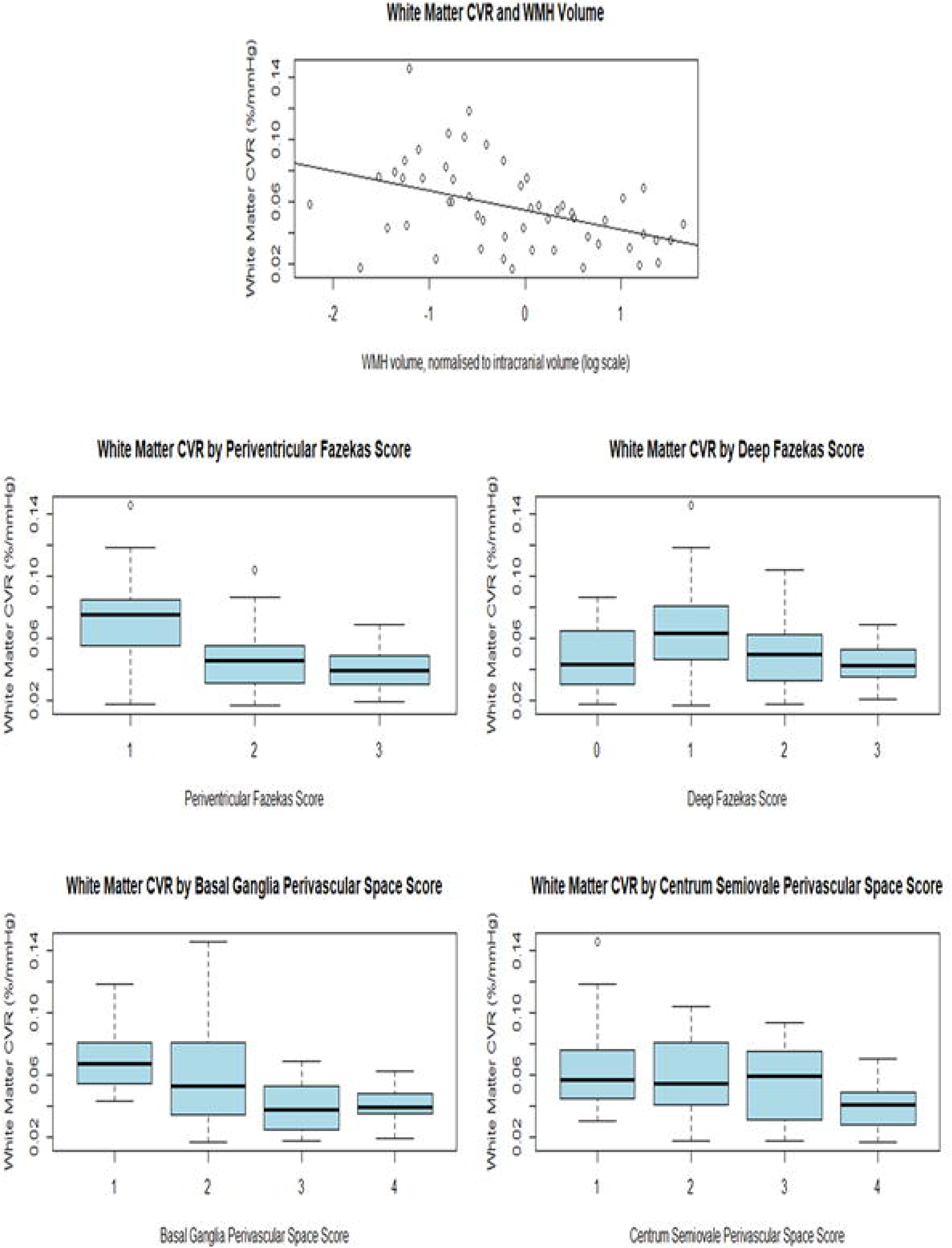
White matter **CVR** and WMH volume, WMH Fazekas score and **PVS** scores.

#### CVR, Pulsatility and CBF

Reduced white matter CVR was associated with increased PI and RI in the superior sagittal, straight and transverse sinuses (all p<0.05), but not with ICA or internal jugular vein PI or RI, or with CBF, (Figure 2, Supplementary Table 2).

**Figure 2:**
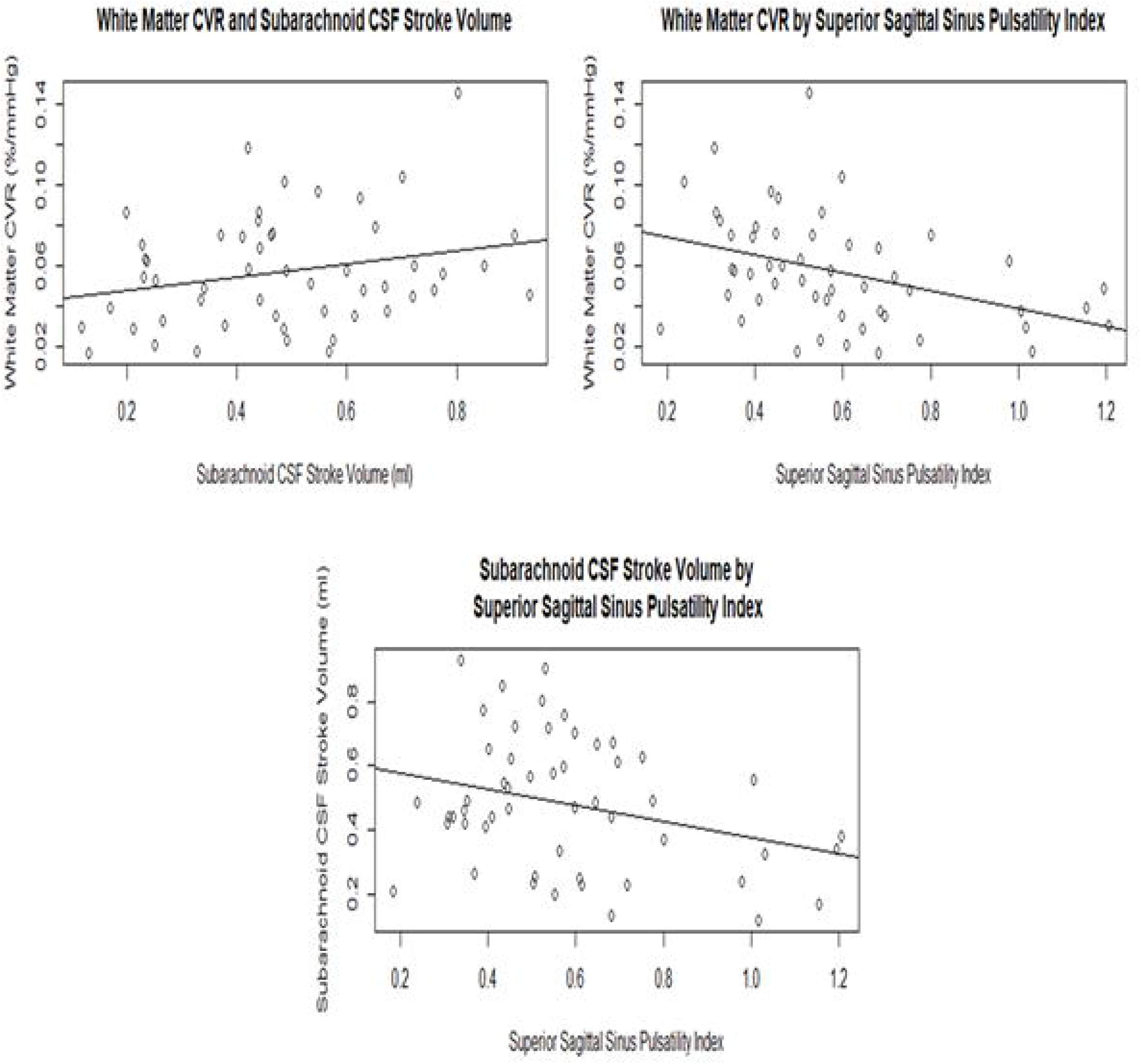
Relationship of white matter CVR, subarachnoid CSF stroke volume and superior sagittal sinus pulsatility index

We found no univariate associations between CSF pulsatility or flow measures and CVR.

#### Co-variate-adjusted analyses

##### CVR and SVD Features

After controlling for age, systolic BP and gender, we found reduced white matter CVR remained associated with WMH volume (−0.01 %/mmHg change in EtCO2 per log10 increase in WMH volume, p=0.02), periventricular WMH Fazekas score (−0.01 %/mmHg change in EtCO2 per point increase, p=0.02) and basal ganglia PVS score (−0.01 %/mmHg change in EtCO2 per point increase, p=0.02). The associations between these SVD features and grey matter CVR was no longer significant (Table 2). We saw a similar pattern when pulse pressure was substituted for systolic BP (data not shown).

Deep and total Fazekas scores, centrum semiovale PVS, lacunes, any brain atrophy, and total SVD score were not associated with reduced CVR in either white or grey matter (Table 2).

We tested the independence of the basal ganglia PVS score association with reduced white matter CVR further by adding WMH volume to the regression model, which resulted in the association no longer being significant (Supplementary Table 3).

**Table 3:**
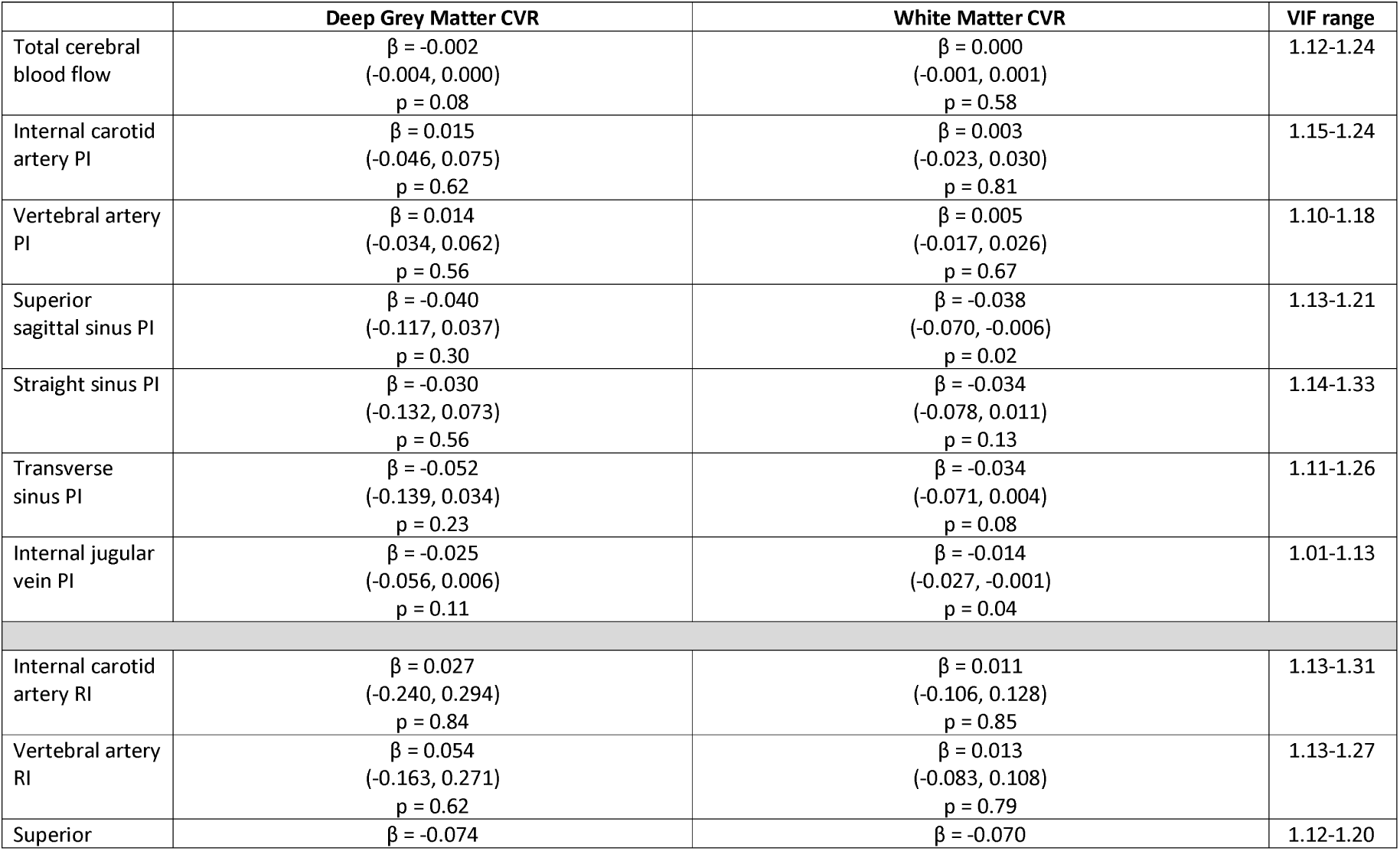
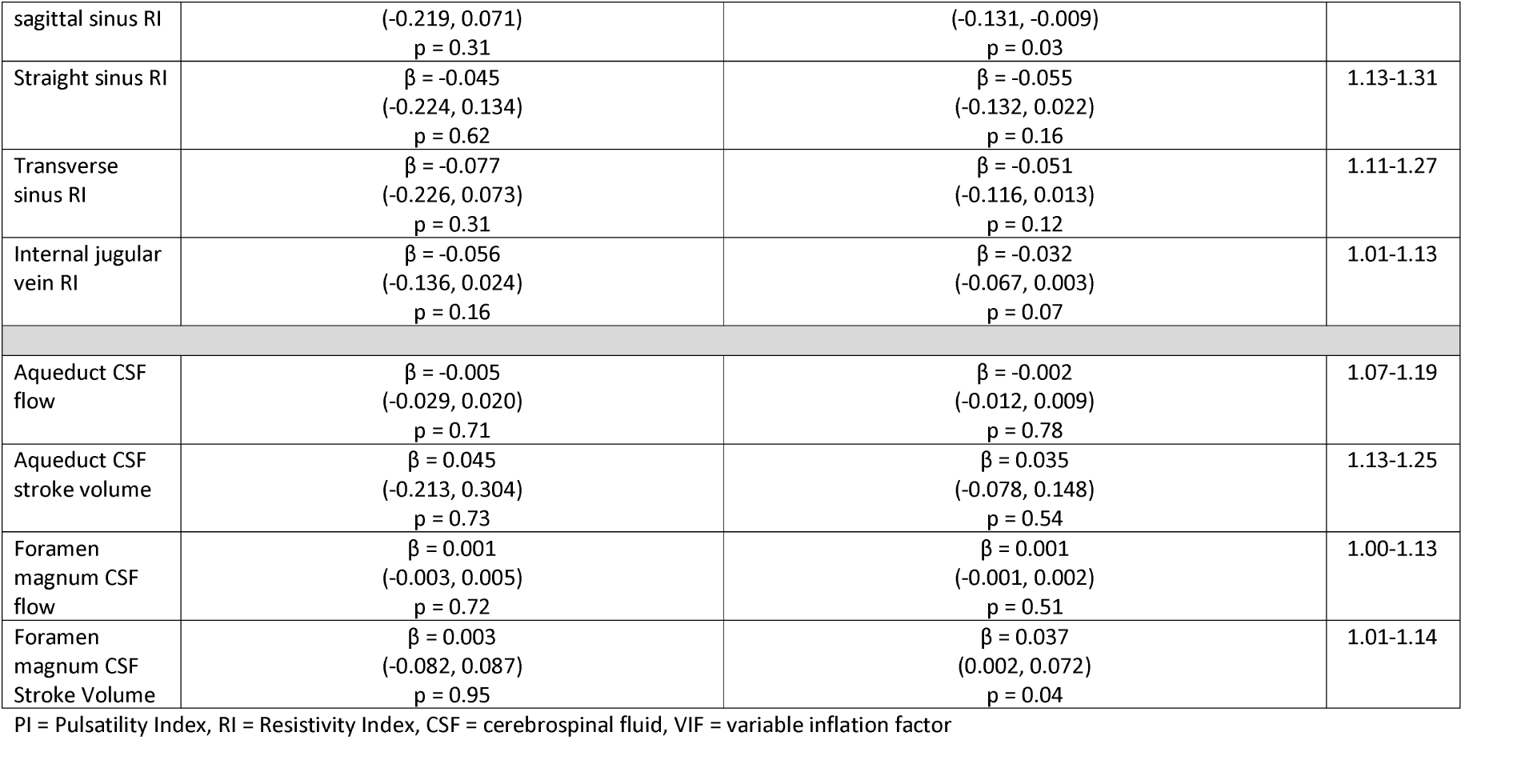
CVR association with CBF, arterial and venous pulsatility and CSF flow dynamics. All multivariate regression models adjusted for age, sex and systolic BP. Standardised β co-efficient, 95% confidence interval and p value.

##### CVR, Pulsatility and CBF

After correcting for age, SBP and gender, reduced white matter CVR remained associated with increasing superior sagittal sinus PI (−0.03 %/mmHg change in EtCO2 per unit increase in PI p=0.02), superior sagittal sinus RI (−0.07%/mmHg change in EtCO2 per unit increase in RI, p=0.03), and internal jugular vein PI (−0.01%/mmHg change in EtCO2 per unit increase in PI, p=0.04), Table 3.

Reduced white matter CVR was also associated with reduced CSF stroke volume at the foramen magnum (−0.04%/mmHg change in EtCO2 per ml decrease in CSF stroke volume, p=0.04). Furthermore, reduced CSF stroke volume was associated with increased basal ganglia PVS score (- 0.96ml per point increase in PVS score, p=0.09, Supplementary Table 4). We saw a similar pattern when pulse pressure was substituted for systolic BP (data not shown).

There was no association of white or grey matter CVR with arterial PI or RI, or with CBF (Table 3).

We further tested the association of superior sagittal sinus and internal jugular vein pulsatility metrics by adding WMH volume into the regression model. Internal jugular vein (but not SSS) PI remained associated with reduced white matter CVR (−0.01%/mmHg change in EtCO2 per unit increase in PI, p<0.05, Supplementary Table 3).

## Discussion

We measured concurrently several indices of intracranial microvascular function in patients with minor ischaemic stroke and features of SVD on neuroimaging. We demonstrate that reduced CVR is associated with increased WMH burden and increased basal ganglia PVS, independently of age, gender and blood pressure. The association was stronger for white matter than for subcortical grey matter CVR. Furthermore, the association between reduced CVR and WMH was stronger for periventricular than deep WMH, perhaps reflecting more periventricular vulnerability to haemodynamic changes at distal end arteries, or differing pathogeneses of WMH by brain region, providing in vivo support in humans for two longstanding hypotheses. We also demonstrated that reduction in white matter CVR was associated with increased intracranial vascular pulsatility, most clearly seen in the intracranial venous sinuses and internal jugular veins, and with increased PVS visibility.(23) Importantly, we also found that reduced CVR was associated with reduced CSF stroke volume at the foramen magnum, and also that reduced CSF stroke volume was associated with increased basal ganglia PVS, showing for the first time in the intact human cranium, a possible link between dysfunctional brain interstitial fluid drainage (evidenced by increased PVS visibility) and impaired CSF pulsation (thought to help flush interstitial fluid through the glymphatic system).(30) We saw no association of reduced CVR in white or grey matter with resting CBF, or of resting CBF with any measures of SVD.

We recruited patients representative of the range of SVD features present in patients with typical lacunar or minor cortical ischaemic stroke to ensure relevance to patients that are commonly affected by SVD.(31, 32) Confirming that vascular function changes in SVD are not confined to one stroke type increases the generalisability of our findings. Stroke type and SVD imaging features were carefully assessed by experienced specialists using standardised, validated image processing and analysis techniques. CVR was measured by a robust and standardised technique with quantified reliability and reproducibility.(11) We were careful to control the statistical analyses for key patient characteristics and WMH volume where appropriate, all guided by a professional statistician.

There are limitations. Whilst this study is one of the largest in the literature to assess CVR in SVD(12) and the only study so far in humans to assess CVR, vascular and CSF pulsatility and CBF simultaneously, the sample size limits the number of comparisons and adjustment variables. We used a 1.5T MRI scanner, however the main impact of this versus a 3T scanner is less ‘crisp’ structural resolution of the BOLD MRI image which may affect the precision of tissue localisation, but not signal magnitude.(33) However, since we registered all the images into a common image processing space and used ROIs obtained from high quality structural images mapped onto the CVR image to extract the tissue-specific CVR signals, the 1.5T MRI is unlikely to have affected the tissue associations.

We found a strong association of reduced CVR with WMH visual score and volume. However, the Fazekas score allowed us to detect the novel finding of a stronger association between reduced CVR and periventricular than deep WMH. This could reflect reduced vascularity of the distal perforating arterioles and is consistent with recent findings in two similar populations where reduced retinal arteriolar branching complexity (the retina being a surrogate for brain microvessels) was associated with WMH and other SVD features.(34) The finding of a stronger association between reduced CVR and periventricular rather than deep WMH should be replicated in future studies which should examine regional WMH rather than just total volume, since we show that total volume may obscure important differences in relationships between SVD lesions and microvascular function.

Previous studies found various associations between CVR and WMH in different populations. Yezhuvath et al (n=25) found reduced CVR with increasing WMH volume in Alzheimer’ s disease.(35) Hund-Georgiadis et al (n=17) found reduced CVR in association with a composite SVD severity score including WMH in patients with stroke.(36) Uh et al (n=33) found that CVR in normal white matter declined with increasing WMH volume in older participants with WMH.(37) Van Opstal et al. (n=60) demonstrated BOLD reactivity in the occipital cortex is reduced in individuals with symptomatic hereditary cerebral amyloid angiopathy who had severe WMH and microbleeds compared to controls and presymptomatic carriers.(38)

In several papers by Sam et al. a cohort of patients (n=45-75) over 50 years of age with WMH of Fazekas grade 2 or more, presenting to a neurology clinic with a wide variety of symptoms, demonstrated several associations with altered CVR. Areas of normal white matter with both lower and delayed CVR progressed to become WMH(13, 15) and areas with negative CVR (reduced blood flow below baseline during CO2 inhalation, possibly suggestive of a vascular steal phenomenon) were associated with lower CBF and microstructural damage on DTI.(14) Lower CVR, FA, CBF and increased T2 and MD were observed in WMH compared with contralateral normal white matter.(16)

In contrast, three studies did not show associations between CVR and WMH, although WMH burden was low in all three. Gauthier et al (n=85) found no association of WMH volume with CVR, however the mean WMH volume in their cohort was small (1.0±1.7ml)(39) and they did find CVR in periventricular watershed areas to be positively associated with general fitness assessed as maximum aerobic capacity (VO_2_ max).(39) Richiardi et al. (n=63) found no association between their unique “CVR velocity” metric and Fazekas score in patients with Alzheimer’s disease or mild cognitive impairment or healthy older subjects(40) but the WMH burden was low and their CVR velocity method is very different to that used here and in most prior studies (CO_2_ delivery via nasal cannulae and no monitoring of EtCO_2_). Conijn et al. (n=49) found no association of CVR and WMH volume in younger subjects with few WMH.(41)

Our systematic review highlighted the variability of CVR associations with regard to patient characteristics.(12) The association of reduced CVR with increased BP was seen in some prior studies (41, 42) but BP associations with CVR were unclear in many other studies. A trend for increasing age to be associated with reduced CVR narrowly failed to reach significance in our study, but age was associated with reduced CVR in several other studies.(36, 39, 41, 43) The lack of a definite age-CVR relationship in the present study may reflect poorer brain vascular health in the younger compared with the older stroke patients since patients who experience a stroke aged in their 40s or 50s may have worse vascular health than those who do not experience stroke until their ninth decade. This paradox has almost certainly dampened the expected age effect.

Reduced CVR may be associated with other markers of vascular health and symptoms. Reduced CVR was associated with raised inflammatory markers(44) and more rapid decline in gait speed in diabetics(45) and with decreased insulin sensitivity in obese individuals with insulin resistance.(46) The literature on CVR and cognitive performance is inconclusive.(47)

We are the first to describe an association of increased PVS with reduced CVR and CSF stroke volume, in addition to increased intracranial vascular pulsatility.(23) Visible PVS are associated with hypertension, increasing age, systemic and intracerebral inflammation, other SVD features like WMH and microbleeds, and increased risk of dementia.(1) Longitudinal studies are required to demonstrate if PVS visibility precedes or follows impairments in CVR or changes in vascular or CSF pulsatility. PVS are part of the brain’s waste clearance and immune defence system, recently referred to as the glymphatic system;(48) in this regard, it is interesting that we detected altered pulsation of the CSF at the foramen magnum in association with reduced CVR and with worse PVS since, in experimental models, reduced PVS flushing may lead to failed clearance of metabolic debris and PVS dilation.(30, 48) CSF stroke volume at the foramen magnum was recently shown to be related to inspiration and improved venous return(49) – we extend this work by demonstrating relationships between CSF stroke volume and increased PVS visibility. Reduced CSF stroke volume could indicate reduced CSF flux around the base of the brain with reduced CSF movement linked to impaired PVS flushing. Caution is required, however, since the PVS-CSF stroke volume and CVR associations could be a co-association with other SVD features, a common problem in SVD research(6) and should be evaluated further.

We did not find an association between WMH and CBF, consistent with prior data.(4) However, we provide more support for a vascular dysfunction in SVD: CVR is a measure of dynamic vessel function, with impairment reflecting an inability to increase blood flow when required. The mechanisms behind this may encompass impaired endothelial function, reduced nitric oxide bioavailability, altered vascular smooth muscle function, inflammatory exudates damaging the vessel wall and perivascular space function.(50) More longitudinal data with contemporaneous vascular function measures and careful adjustment for co-variates are required to unpick this pathway, but it is notable that all the associations of demographic and SVD features with impaired CVR were stronger in the white matter than the grey matter despite white matter CVR being around one third the magnitude of that in grey matter and white matter CBF being around half of that in grey matter.(4, 23) Studies of SVD mechanisms and potential interventions should consider using measures of dynamic vascular function such as CVR or pulsatility rather than resting CBF.(5, 38)

### Conclusions and future directions

We show novel and independent associations between SVD features and several measures of impaired cerebral haemodynamics, providing new insights into mechanisms that may underlie SVD development. Further longitudinal studies, controlling for confounders (age, BP, WMH volume, etc) will define these changes and delineate the pathway of SVD development, including the novel observation of impaired CSF pulsation. Ultimately, enhanced knowledge of vascular malfunction will help identify therapeutic targets to halt or even reverse disease progression, with benefits for both dementia and stroke prevention.

