## Supplemental tables for "Intracranial functional haemodynamic relationships in patients with cerebral small vessel disease"

**Supplementary Table 1:**

**MRI Imaging Parameters**

| scan | sequence | TR (ms) | TE (ms) | FA (°) | FOV (mm) | Acquisition matrix | slices × thickness (mm) | Other |
| --- | --- | --- | --- | --- | --- | --- | --- | --- |
| axial BOLD CVR | 2D single shot spin-echo EPI | 3000 | 45 | 90 | 256 x 256 | 64 x 64 | 36 x 4 | 8 dummy volumes acquired prior to 12-min paradigm |
| sagittal T1w volume | 3D IR-prepared spoiled gradient echo | 9.6 | 4.0 | 8 | 256 x 256 | 192 x 192 | 160 x 1.3 | TI = 500 ms |
| axial FLAIR | 2D IR-prepared fast spin-echo | 8000 | 100 | 90 | 240 x 240 | 320 x 256 | 36 x 4 | TI = 2000 ms |
| axial T2w | 2D PROPELLER | 7000 | 90 | 90 | 240 x 240 | 384 | 36 x 4 | 1.5 averages |
| axial GRE | 2D gradient echo | 900 | 15 | 20 | 240 x 240 | 384 (AP) x 256 (LR) | 36 x 4 |  |
| aqueduct CSF flow | 2D phase-contrast | 11 | 6.5 | 20 | 160 x 160 | 256 x 256 | 1 x 6 | 2 averages, 32 phases, 2 views per segment, venc = 10 cm/s |
| venous sinuses flow (coronal-oblique) | 2D phase-contrast | 9 | 5 | 25 | 160 x 160 | 256 (SI) x 128 (LR) | 1 x 6 | 2 averages, 32 phases, 2 views per segment, venc = 50 cm/s |
| axial blood flow (neck) | 2D phase-contrast | 9 | 5 | 25 | 160 x 160 | 256 (AP) x 128 (LR) | 1 x 6 | 2 averages, 32 phases, 2 views per segment, venc = 70 cm/s |
| axial cervical CSF flow | 2D phase-contrast | 11 | 7 | 20 | 160 x 160 | 256 (AP) x 128 (LR) | 1 x 6 | 3 averages, 32 phases, 2 views per segment, venc = 6 cm/s |

**Supplementary Table 2: Univariate Linear regression of CVR with baseline demographic features, SVD features, CBF, vascular pulsatility and CSF dynamic measures.** Standardised β co-efficient, 95% confidence interval and p value.

|  | **Deep Grey Matter CVR** | **White Matter CVR** |
| --- | --- | --- |
| Age | β = -0.000  (-0.002, 0.002)  p = 0.88 | β = -0.001  (-0.002, 0.000)  p = 0.09 |
| Sex | β = 0.005  (-0.034, 0.045)  p = 0.79 | β = 0.000  (-0.017, 0.018)  p = 0.98 |
| NIHSS | β = 0.005  (-0.010, 0.019)  p = 0.53 | β = 0.002  (-0.004, 0.009)  p = 0.47 |
| Rankin | β = 0.014  (-0.023, 0.050)  p = 0.45 | β = 0.008  (-0.009, 0.024)  p = 0.36 |
| History of TIA | β = -0.028  (-0.155, 0.099)  p = 0.66 | β = 0.002  (-0.055, 0.058)  p = 0.96 |
| History of Stroke | β = 0.023  (-0.028, 0.074)  p = 0.37 | β = -0.004  (-0.027, 0.019)  p = 0.72 |
| History of Peripheral Vascular Disease | β = 0.032  (-0.095, 0.159)  p = 0.62 | β = 0.007  (-0.050, 0.063)  p = 0.82 |
| History of Diabetes | β = -0.032  (-0.083, 0.018)  p = 0.20 | β = -0.020  (-0.042, 0.003)  p = 0.08 |
| History of Hypertension | β = 0.001  (-0.039, 0.041)  p = 0.96 | β = 0.003  (-0.015, 0.021)  p = 0.76 |
| Antihypertensive use Pre-Stroke | β = 0.001  (-0.034, 0.036)  p = 0.95 | β = -0.003  (-0.019, 0.012)  p = 0.68 |
| History of Atrial Fibrillation | β = -0.023  (-0.082, 0.036)  p = 0.44 | β = -0.029  (-0.055, -0.004)  p = 0.02 |
| Statin Use at Time of Stroke | β = -0.004  (-0.040, 0.032)  p = 0.84 | β = -0.013  (-0.029, 0.003)  p = 0.11 |
| History of Hyperlipidaemia | β = 0.001  (-0.035, 0.038)  p = 0.94 | β = -0.003  (-0.019, 0.014)  p = 0.73 |
| History of Structural Heart Disease | β = -0.018  (-0.097, 0.061)  p = 0.64 | β = -0.022  (-0.056, 0.013)  p = 0.22 |
| Family History of Stroke | β = 0.027  (-0.015, 0.069)  p = 0.20 | β = 0.019  (0.001, 0.038)  p = 0.04 |
| History of Any Ischaemic Heart Disease | β = 0.012  (-0.047, 0.072)  p = 0.68 | β = -0.004  (-0.031, 0.022)  p = 0.74 |
| Smoking History – Current vs Never | β = -0.020  (-0.057, 0.017)  p = 0.29 | β = 0.004  (-0.013, 0.021)  p = 0.64 |
| Smoking History – Current or ex-smokerless than one year vs Never or ex-smoker more than one year | β = -0.024  (-0.063, 0.016)  p = 0.24 | β = 0.004  (-0.014, 0.022)  p = 0.69 |
| Smoking History – Any history vs never | β = -0.005  (-0.041, 0.03)  p = 0.76 | β = 0.004  (-0.012, 0.019)  p = 0.66 |
| History of Alcohol Excess | β = -0.001  (-0.046, 0.043)  p = 0.94 | β = 0.007  (-0.012, 0.027)  p = 0.46 |
| Alcohol consumption (per unit increase) | β = -0.001  (-0.001, 0.001)  p = 0.31 | β = -0.000  (-0.000, 0.000)  p = 0.52 |
| Current antiplatelet use | β = 0.020  (-0.039, 0.079)  p = 0.50 | β = 0.026  (0, 0.051)  p = 0.05 |
| Current anticoagulant use | β = -0.032  (-0.091, 0.026)  p = 0.27 | β = -0.028  (-0.053, -0.003)  p = 0.03 |
| Current statin use | β = -0.031  (-0.089, 0.028)  p = 0.30 | β = -0.010  (-0.036, 0.017)  p = 0.46 |
| Systolic Blood Pressure | β = -0.001  (-0.002, -0.000)  p = 0.01 | β = -0.001  (-0.001, -0.000)  p = 0.01 |
| Diastolic Blood Pressure | β = -0.001  (-0.003, 0.001)  p = 0.36 | β = -0.000  (-0.001, 0.001)  p = 0.47 |
| Mean Arterial Pressure | β = -0.002  (-0.004, 0.000)  p = 0.06 | β = -0.001  (-0.002, 0.000)  p = 0.08 |
| Pulse Pressure | β = -0.002  (-0.003, -0.000)  p = 0.01 | β = -0.001  (-0.001, -0.000)  p = 0.01 |
| White Matter Hyperintensity Volume (log scale) | β = -0.017  (-0.035, 0.001)  p = 0.06 | β = -0.013  (-0.020, -0.005)  p = <0.01 |
| Fazekas Score - Deep WMH | β = -0.011  (-0.031, 0.009)  p = 0.29 | β = -0.008  (-0.017, 0.001)  p = 0.08 |
| Fazekas Score Periventricular WMH | β = -0.023  (-0.044, -0.003)  p = 0.03 | β = -0.015  (-0.024, -0.007)  p = <0.01 |
| Fazekas Score – Total Score | β = -0.009  (-0.020, 0.001)  p = 0.09 | β = -0.006  (-0.011, -0.002)  p = 0.01 |
| PVS Score – Basal Ganglia | β = -0.018  (-0.036, -0.000)  p = 0.05 | β = -0.012  (-0.020, -0.005)  p = <0.01 |
| PVS Score – Centrum Semiovale | β = -0.016  (-0.034, 0.001)  p = 0.07 | β = -0.009  (-0.016, -0.001)  p = 0.03 |
| Presence of Lacunes | β = -0.029  (-0.065, 0.006)  p = 0.11 | β = -0.018  (-0.034, -0.003)  p = 0.02 |
| Brain parenchymal volume (normalised to intracranial volume) | β = 0.249  (-0.281, 0.780)  p = 0.35 | β = 0.061  (-0.178, 0.299)  p = 0.61 |
| Superficial Atrophy | β = -0.010  (-0.049, 0.029)  p = 0.59 | β = -0.011  (-0.028, 0.006)  p = 0.21 |
| Deep Atrophy | β = -0.014  (-0.046, 0.019)  p = 0.39 | β = -0.015  (-0.029, -0.001)  p = 0.04 |
| Stroke Type – Lacunar vs Cortical | β = 0.003  (-0.035, 0.040)  p = 0.89 | β = 0.000  (-0.016, 0.017)  p = 0.97 |
| Total SVD Score | β = -0.01  (-0.023, 0.003)  p = 0.12 | β = -0.007  (-0.013, -0.002)  p = 0.01 |
| Total CBF | β = -0.002  (-0.003, 0.000)  p = 0.06 | β = -0.000  (-0.001, 0.001)  p = 0.73 |
| ICA PI | β = -0.008  (-0.066, 0.050)  p = 0.78 | β = -0.009  (-0.034, 0.016)  p = 0.48 |
| VA PI | β = 0.001  (-0.047, 0.048)  p = 0.98 | β = -0.003  (-0.024, 0.018)  p = 0.75 |
| SSS PI | β = -0.058  (-0.129, 0.013)  p = 0.11 | β = -0.044  (-0.073, -0.014)  p = <0.01 |
| StS PI | β = -0.058  (-0.149, 0.033)  p = 0.20 | β = -0.046  (-0.085, -0.008)  p = 0.02 |
| TS PI | β = -0.078  (-0.159, 0.002)  p = 0.06 | β = -0.044  (-0.078, -0.010)  p = 0.01 |
| IJV PI | β = -0.024  (-0.056, 0.009)  p = 0.15 | β = -0.013  (-0.027, 0.001)  p = 0.06 |
| ICA RI | β = -0.08  (-0.330, 0.152)  p = 0.46 | β = -0.053  (-0.158, 0.052)  p = 0.32 |
| VA RI | β = -0.039  (-0.242, 0.164)  p = 0.70 | β = -0.035  (-0.124, 0.053)  p = 0.43 |
| SSS RI | β = -0.108  (-0.244, 0.03)  p = 0.11 | β = -0.082  (-0.138, -0.026)  p = 0.01 |
| StS RI | β = -0.100  (-0.259, 0.059)  p = 0.21 | β = -0.079  (-0.147, -0.012)  p = 0.02 |
| TS RI | β = -0.126  (-0.265, 0.013)  p = 0.07 | β = -0.071  (-0.131, -0.011)  p = 0.02 |
| IJV RI | β = -0.055  (-0.137, 0.028)  p = 0.19 | β = -0.030  (-0.066, 0.006)  p = 0.10 |
| Aqueduct flow | β = -0.007  (-0.031, 0.017)  p = 0.55 | β = -0.002  (-0.013, 0.008)  p = 0.68 |
| Aqueduct stroke volume | β = 0.066  (-0.184, 0.316)  p = 0.60 | β = 0.024  (-0.086, 0.134)  p = 0.66 |
| Foramen magnum CSF flow | β = 0.001  (-0.003, 0.005)  p = 0.74 | β = 0.001  (-0.001, 0.002)  p = 0.54 |
| Foramen magnum Stroke Volume | β = -0.008  (-0.095, 0.078)  p = 0.85 | β = 0.033  (-0.004, 0.070)  p = 0.08 |

**Supplementary Table 3. Multivariate regression models from Table 2/3 additionally adjusted for WHM volume. All models adjusted for age, sex, systolic blood pressure and WMH volume.**

|  | **Deep Grey Matter CVR** | **White Matter CVR** | **VIF range** |
| --- | --- | --- | --- |
| PVS Score – Basal Ganglia | β = -0.014  (-0.035, 0.008)  p = 0.20 | β = -0.008  (-0.017, 0.002)  p = 0.10 | 1.10-1.77 |
| PVS Score – Centrum semiovale | β = -0.013  (-0.031, 0.005)  p = 0.17 | β = -0.005  (-0.013, 0.003)  p = 0.22 | 1.12-1.71 |
| SSS PI | β = -0.018  (-0.098, 0.061)  p = 0.65 | β = -0.028  (-0.061, 0.005)  p = 0.10 | 1.18-1.66 |
| SSS RI | β = -0.036  (-0.184, 0.112)  p = 0.63 | β = -0.052  (-0.114, 0.010)  p = 0.10 | 1.16-1.63 |
| IJV PI | β = -0.023  (-0.054, 0.007)  p = 0.13 | β = -0.013  (-0.026, -0.000)  p = <0.05 | 1.01-1.53 |
| Subarachnoid Stroke Volume | -0.015  (-0.099, 0.069)  p = 0.72 | 0.030  (-0.006, 0.064)  p = 0.10 | 1.13-1.56 |

**Supplementary Table 4: Association of CSF flow at the cerebral aqueduct and foramen magnum with PVS score. Models adjusted for age, sex and systolic blood pressure.** Standardised β co-efficient, 95% confidence interval and p value.

|  | **Basal Ganglia PVS Score** | **Centrum Semiovale PVS Score** |
| --- | --- | --- |
| Cerebral Aqueduct CSF Stroke Volume | β = -0.841  (-4.367 , 2.685)  p = 0.63 | β = 0.200  (-3.891 , 4.291)  p = 0.92 |
| Foramen Magnum CSF Stroke Volume | β = -0.971  (-2.087 , 0.146)  p = 0.09 | β = -0.275  (-1.607 , 1.056)  p = 0.68 |
